# The transmembrane protein Syndecan regulates stem cell nuclear properties and cell maintenance

**DOI:** 10.1101/2024.02.15.580237

**Authors:** Buffy L. Eldridge-Thomas, Jerome G. Bohere, Chantal Roubinet, Alexandre Barthelemy, Tamsin J. Samuels, Felipe Karam Teixeira, Golnar Kolahgar

**Affiliations:** Department of Physiology, Development and Neuroscience, University of Cambridge, Cambridge, CB2 3DY, UK; IGDR (Institut de Génétique et Développement de Rennes) – CNRS UMR 6290, 35000, Rennes, France; Present address: Institut Curie, Laboratory of Genetics and Developmental Biology, PSL Research University, INSERM U934, CNRS UMR3215, 75005 Paris, France; Department of Genetics, University of Cambridge, Cambridge, CB2 3EH, UK

## Abstract

Tissue maintenance is underpinned by resident stem cells whose activity is modulated by microenvironmental cues. Using *Drosophila* as a simple model to identify regulators of stem cell behaviour and survival *in vivo*, we have identified novel connections between the conserved transmembrane proteoglycan Syndecan, nuclear properties and stem cell function. In the *Drosophila* midgut, Syndecan depletion in intestinal stem cells results in their loss from the tissue, impairing tissue renewal. At the cellular level, Syndecan depletion alters cell and nuclear shape, and causes nuclear lamina invaginations and DNA damage. In a second tissue, the developing *Drosophila* brain, live imaging revealed that Syndecan depletion in neural stem cells results in nuclear envelope remodelling defects which arise upon cell division. Our findings reveal a new role for Syndecan in the maintenance of nuclear properties in diverse stem cell types.

**Graphical Abstract:** 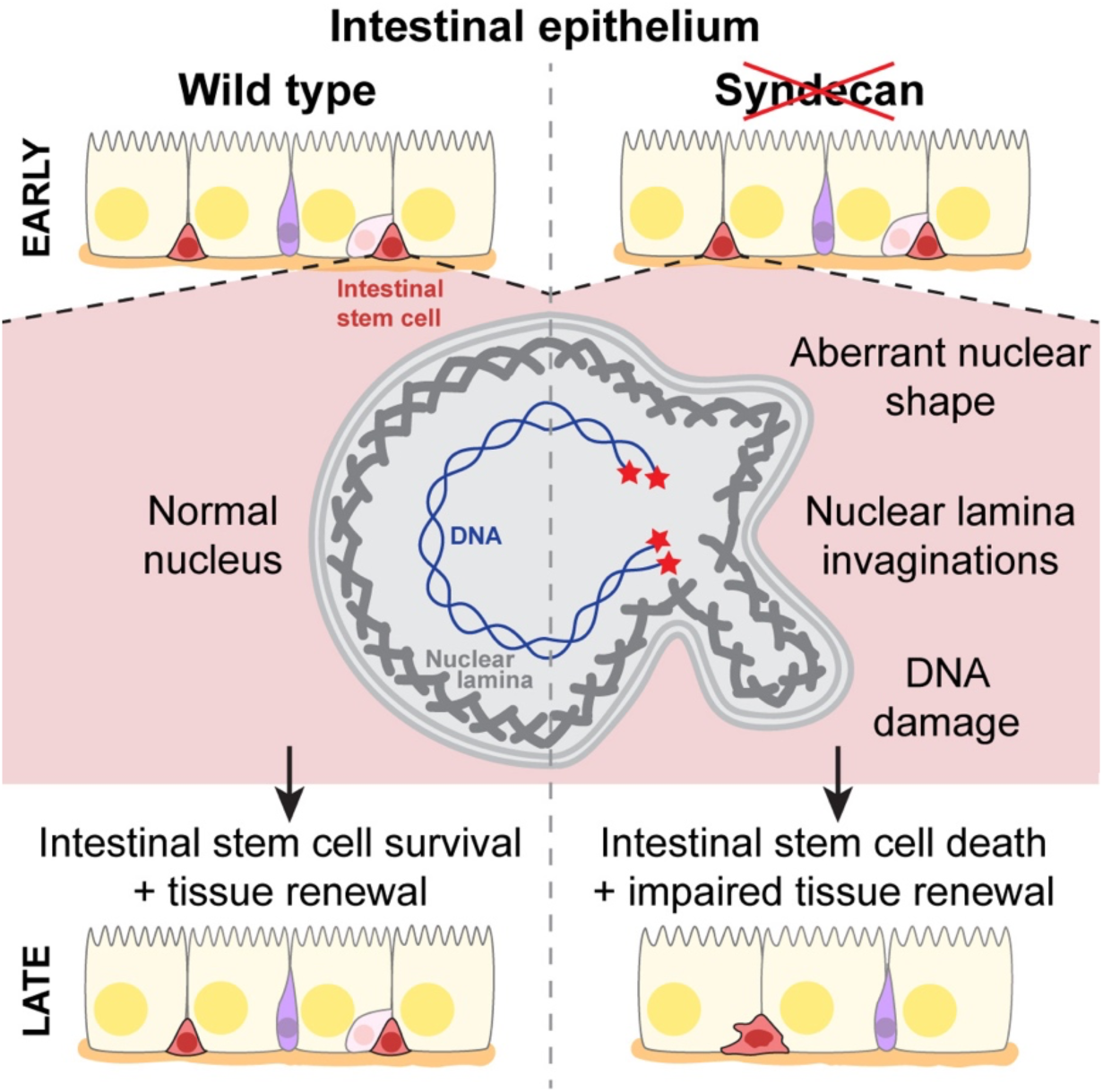

## Introduction

Most tissues are renewed by resident stem cells, capable of producing new specialised cells when required. Mechanisms promoting stem cell maintenance include genome protection, induced quiescence and stem cell self-renewal through asymmetric cell divisions (Beumer & Clevers, 2024). Deregulation of these mechanisms can lead to acquisition of mutations and cancer development or stem cell loss and tissue attrition (Beumer & Clevers, 2024).

Stem cell survival and fate decisions are influenced by the basement membrane, a specialised extracellular matrix whose complex structure and composition varies dynamically with physiological context (Jayadev & Sherwood, 2017; Pozzi et al., 2017; Sekiguchi & Yamada, 2018). Whilst *in vitro* systems are often used to study stem cell behaviour, they incompletely recapitulate this extracellular environment, highlighting the ongoing value of *in vivo* models to further our understanding of stem cell biology and disease. To investigate stem cells in their native milieu, we turned to the simple, genetically tractable *Drosophila*. The adult *Drosophila* gut is maintained throughout life by resident stem cells (Micchelli & Perrimon, 2006; Ohlstein & Spradling, 2006) (Figure 1A-A’), allowing long term assessment of stem cell maintenance and tissue-level effects, while the developing larval brain contains rapidly dividing neural stem cells highly amenable to live imaging, facilitating a more dynamic understanding of cell biological events. While testing the role of basement membrane receptors in stem cell activity, we investigated the single *Drosophila* ortholog of Syndecan (Sdc) (Spring et al., 1994).

**Figure 1.**
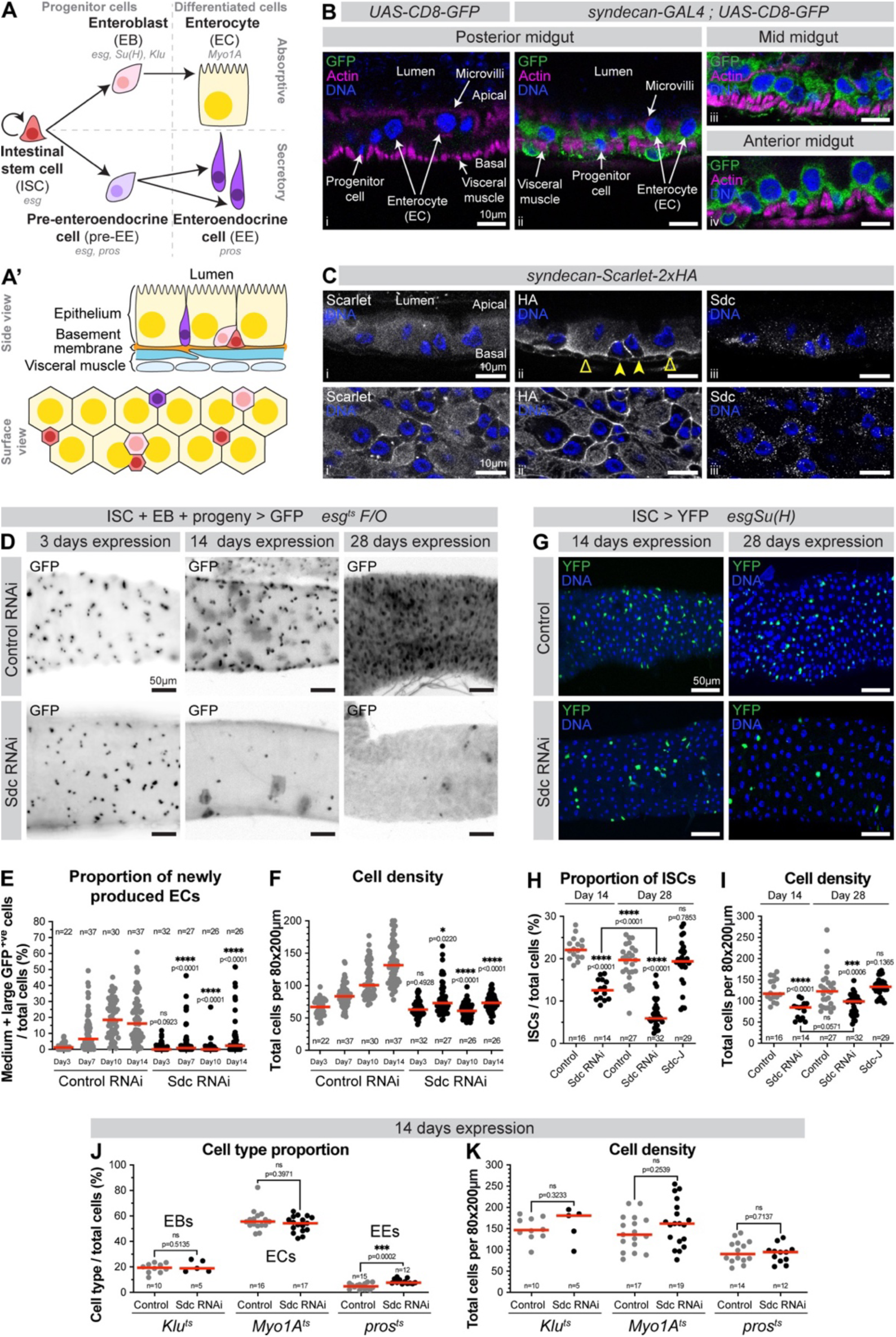
Sdc is required for *Drosophila* intestinal stem cell maintenance. (A) Intestinal cell lineages. Intestinal stem cells (ISCs) self-renew and give rise to enteroblasts (EBs) which differentiate into absorptive enterocytes (ECs); and pre-enteroendocrine cells (pre-EEs) which undergo one division before differentiating into a pair of secretory enteroendocrine cells. Progenitor cells comprise ISCs, EBs and pre-EEs. Cell type-specific genes shown in italics. (A’) Side and surface view schematics of the posterior midgut. (B) Side views of midguts carrying either *UAS-CD8-GFP* alone (i) or with *syndecan-GAL4* (ii-iv). Progenitor cells are identified by their small, basally located nuclei. ECs are identified by their large nuclei. DNA stain (blue) marks nuclei; GFP (green) reports *sdc* expression; Phalloidin (magenta) marks the visceral muscle and EC microvilli. (C) Side (i-iii) and basolateral surface views (i’-iii’) of midguts expressing endogenous Sdc tagged with Scarlet and HA at its C-terminus. ECs (empty arrowheads) show diffuse, basal Sdc signal. Progenitors/EEs (filled arrowheads) show Sdc enrichment. DNA stain (blue) marks nuclei; Scarlet, anti-HA and anti-Sdc (all white) mark Sdc protein. (D) Surface views of midguts expressing control RNAi or Sdc RNAi using the *esg^ts^ F/O* system. GFP (black) marks small progenitor cells and their progeny. (E) Proportion of medium and large GFP^+ve^ cells (corresponding to newly produced differentiating and terminally differentiated ECs, respectively), and (F) Cell density. n=number of guts, from three replicates. (G) Surface views of midguts expressing Sdc RNAi in ISCs using the *esgSu(H)* system. DNA stain (blue) marks the nuclei of all cells; YFP (green) marks ISCs. (H) ISC proportion and (I) Cell density. n=number of guts, from two (Day 14) or three (Day 28) replicates. (J) Cell type proportion and (K) Cell density of midguts expressing Sdc RNAi in EBs (*Klu^ts^*), ECs (*Myo1A^ts^*) or pre-EEs and EEs (*pros^ts^*). n=number of guts, from one (EBs) or three (ECs & EEs) replicates.

Sdc proteins are transmembrane proteoglycans bearing heparan and chondroitin sulfate chains on their extracellular domain, via which they bind to basement membrane components including Laminin and Collagen IV (Xian et al., 2010; Gondelaud & Ricard-Blum, 2019). Mammals possess four *sdc* genes with varying extracellular domains and a highly conserved small cytoplasmic domain involved in protein-protein interactions, including direct and indirect association with the microtubule and actin cytoskeletons (Yoneda & Couchman, 2003; Gondelaud & Ricard-Blum, 2019). Sdc proteins play important and diverse roles during development, inflammation, disease, and tissue repair, through adhesion and signalling (Couchman et al., 2015; Afratis et al., 2017; Ravikumar et al., 2020; Ricard-Blum & Couchman, 2023). However, so far, little is known about Sdc’s role in stem cell maintenance and tissue homeostasis.

Here we report that *Drosophila* Sdc contributes to intestinal stem cell maintenance, through pro-survival mechanisms associated with nuclear properties and genome protection. Furthermore, we identify an additional role in regulating nuclear envelope remodelling during asymmetric division of neural stem cells. Our findings uncover a novel connection between Sdc and nuclear properties in multiple stem cell types.

## Results

### Syndecan is required for long-term *Drosophila* intestinal stem cell maintenance

Sdc, a previously uncharacterised player in the adult *Drosophila* intestine, is expressed throughout the intestinal epithelium, in all cell types (Figure 1B-C). To test whether Sdc contributes to intestinal cell production, we knocked down Sdc by RNAi expression in progenitor cells (Figure 1A) using the *esg^ts^ F/O* (“escargot flip out”) system (Jiang et al., 2009) (Methods). This system also drives heritable GFP expression in progenitor cells and their progeny, allowing assessment of progenitor maintenance and new cell production. Expression of three independent RNAi lines resulted in a lack of cell production (Figure 1D-F and Figure S1A-B), and a progressive loss of progenitor cells (Figure 1D and Figure S1C).

To identify in which cell type(s) of the posterior midgut Sdc is required, we performed RNAi knockdown with cell type-specific GAL4 drivers (Figure 1A and Methods). We monitored the proportion of each cell type to assess whether Sdc is required for its maintenance. In addition, since failure in new cell production is accompanied by low epithelial cell density (Figure 1F), we measured cell density as a proxy for new cell production. Sdc knockdown in ISCs (Figure 1A) resulted in a progressive reduction in ISC proportion (Figure 1G-H). This ISC loss was associated with reduced cell density (Figure 1I), presumably due to a failure in new cell production (Figure 1E). Fly survival was not compromised (Figure S2), consistent with other reports using alternative modes of ISC elimination (Jin et al., 2017; Resende et al., 2017). Knockdown in other cell types did not perturb their proportion, except for EEs which became more abundant (Figure 1J). Cell density was comparable to control guts (Figure 1K), suggesting Sdc is dispensable in other epithelial cell types. We then tested whether Sdc overexpression in ISCs would induce an opposite effect on ISC proportion or cell density, but it did not (Figure 1H-I). Altogether, we have identified Sdc as a previously unrecognised but important regulator of stem cell maintenance in the adult *Drosophila* intestine.

### Sdc-depleted intestinal stem cells are partially lost through apoptosis and display aberrant cell shapes

ISCs can be eliminated from the tissue by differentiation or cell death. Differentiation is ruled out by the reduction in differentiated cells upon expression of Sdc RNAi with the *esg^ts^ F/O* lineage tracing system (Figure 1D-E and Figure S1; compare to control guts where production of large GFP^+ve^ enterocytes is clear). To test whether ISC loss occurs by apoptosis, a common form of programmed cell death, we expressed the inhibitor of apoptosis, DIAP1 (Steller, 2008), in these cells. DIAP1 expression partially rescued (~50%) ISC loss caused by Sdc depletion, and did not affect ISC numbers in control guts (Figure 2A-B). Thus, some ISCs depleted of Sdc are eliminated by apoptosis.

**Figure 2.**
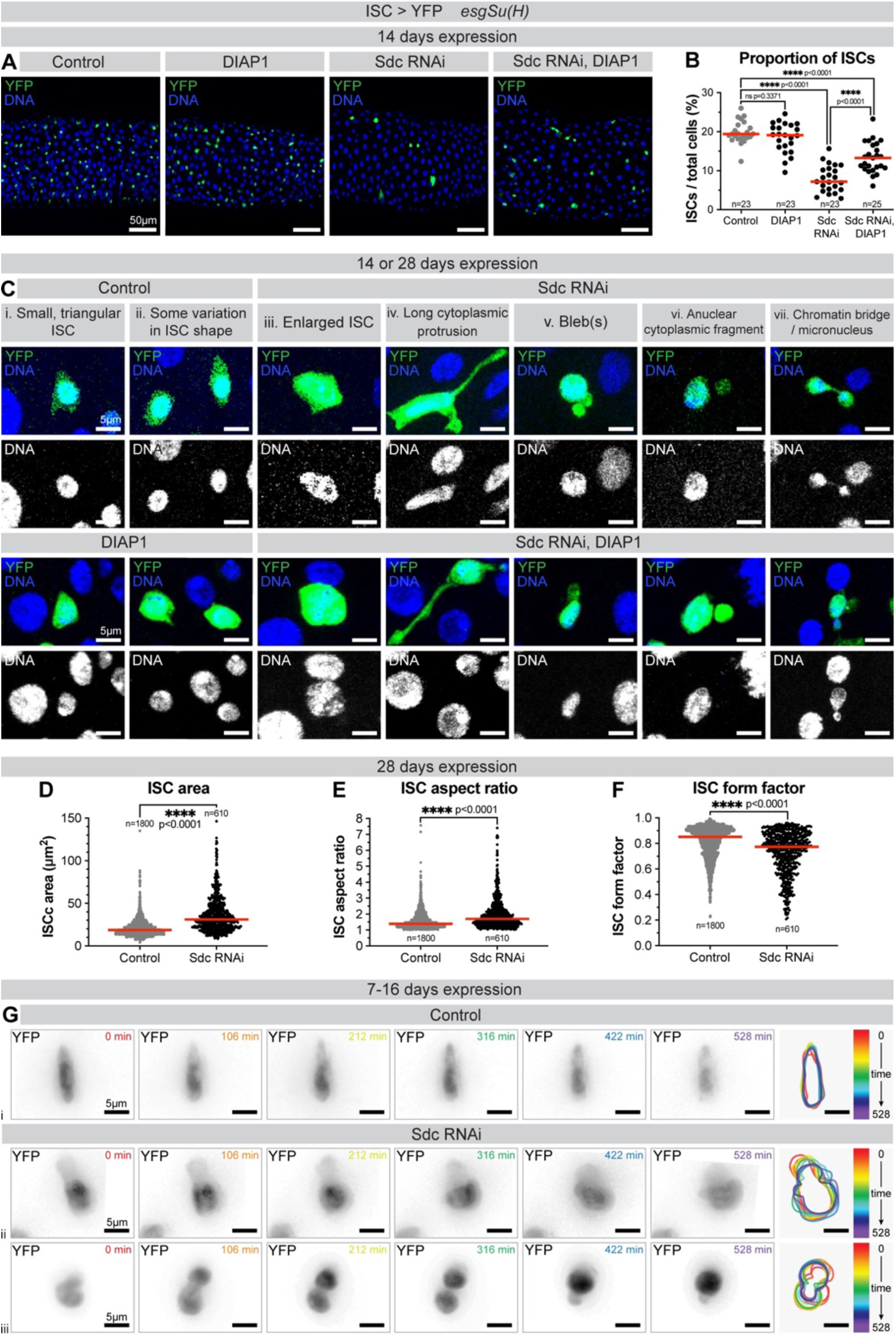
Sdc-depleted ISCs adopt abnormal cell shapes independently of apoptosis. (A) Surface views of midguts expressing +/− DIAP1 and +/− Sdc RNAi in ISCs. DNA stain (blue) marks the nuclei of all cells; YFP (green) marks ISCs. (B) ISC proportion. n=number of guts, from three replicates. (C) Surface views of ISCs. DNA stain (blue/white) marks nuclei; YFP (green) marks ISCs. Images are from either 14 or 28 days expression with *esgSu(H)*. (D) ISC cytoplasmic area; (E) ISC aspect ratio and (F) ISC form factor. n=number of ISCs analysed, from ≥27 guts, from three replicates. (G) Time lapse of control or Sdc RNAi-expressing ISCs. YFP (black) marks ISCs. Right panels show colour-coded snapshots of the cell outline during the time lapse.

We then sought to determine the cellular changes induced by loss of Sdc, which could cause ISC elimination. Observation of labelled Sdc-depleted ISCs at high magnification revealed striking changes in cell shape. *Drosophila* ISCs are normally small and triangular shaped (Ohlstein & Spradling, 2006; Marianes & Spradling, 2013; Hung et al., 2020) (Figure 2Ci-ii). In contrast, Sdc-depleted ISCs showed a variety of morphological defects including larger size, cytoplasmic protrusions, and occasional blebs (Figure 2Ciii-vii). These phenotypes were observed both with and without DIAP1 expression (Figure 2C), indicating that these cell shapes are not caused by apoptosis. Quantification of ISC area and shape revealed that, compared to controls, ISCs expressing Sdc RNAi are significantly larger (Figure 2D), more elongated (Figure 2E) and more convoluted (Figure 2F).

Abnormal cell shape and size might reflect changes in the cortical cytoskeleton and cellular dynamics. We therefore turned to live imaging of fluorescently labelled ISCs in *ex vivo* guts to compare dynamic cellular behaviour over approximately eight hours (>400 ISCs imaged across >14 guts for each genotype). Whilst most control ISCs remained static (Figure 2Gi), Sdc RNAi-expressing ISCs displayed more dynamic protrusions and shape changes (Figure 2Gii). In addition, rare Sdc RNAi-expressing ISCs appeared to attempt but fail in cell division (Figure 2Giii), a behaviour that we have never observed in controls. We concluded that Sdc regulates stem cell shape and that its loss can contribute to changes in cellular morphology compatible with defective divisions.

### Sdc is dispensable for intestinal stem cell abscission but prevents nuclear lamina invaginations and DNA damage

How could Sdc interfere with stem cell morphology and maintenance? Sdc-4 links the abscission machinery to the plasma membrane in mammalian cultured cells, with Sdc-4 depletion delaying abscission or more rarely causing abscission failure, generating binucleate cells (Addi et al., 2020). To explore whether Sdc depletion from proliferating *Drosophila* ISCs impairs their resolution into two separate daughter cells and causes them to become binucleate prior to their elimination, we immunostained guts with Lamin B to label the nuclear compartment and α-catenin to label the cell perimeter (Figure 3A). To confirm that our approach could detect binucleate cells, we used RNAi to knockdown the cytokinesis-promoting Aurora B (Steigemann et al., 2009) (Figure 3Ai-ii). We did not detect any binucleate stem cells upon Sdc knockdown (n>100 ISCs examined across >20 guts) (Figure 3Aiii), indicating that Sdc is not required for ISC abscission.

**Figure 3.**
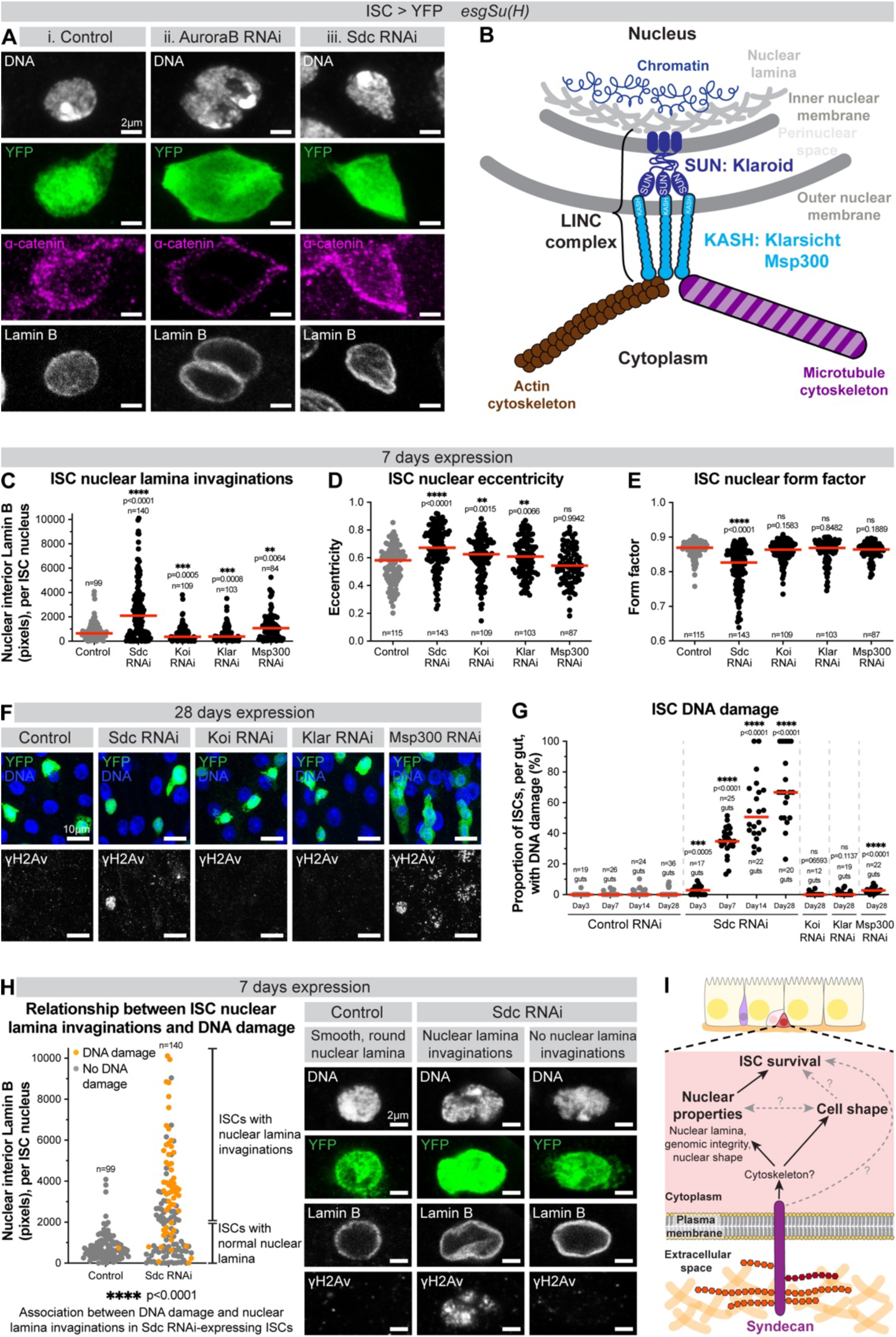
Sdc prevents nuclear lamina invaginations and DNA damage. (A) Control ISC, and ISC expressing Aurora B RNAi or Sdc RNAi. DNA stain (white) marks nuclei; YFP (green) marks ISCs; anti-α-catenin (magenta) marks cell junctions; anti-Lamin B (white) marks nuclear lamina. (B) The *Drosophila* LINC complex. (C) ISC nuclear lamina invaginations (Lamin B pixels located in the nuclear interior), (D) ISC nuclear eccentricity, and (E) ISC nuclear form factor n=number ISCs analysed, from a minimum of 17 guts per genotype, from a minimum of two replicates. (F) Surface view images of midguts. DNA stain (blue) marks nuclei; YFP (green) marks ISCs; anti-γH2Av (white) marks DNA damage. (G) Proportion of ISCs, per gut, with DNA damage. n=number of guts, from a minimum of two replicates. >100 ISCs analysed per genotype per timepoint. (H) Relationship between ISC nuclear lamina invaginations and DNA damage. Each dot represents an individual ISC, coloured orange if ISC has DNA damage and grey if ISC does not. ISCs are defined as having nuclear lamina invaginations if there are ≥2000 Lamin B pixels in the nuclear interior. n=number ISCs analysed, from ≥22 guts per genotype, from four replicates. Example ISCs, with and without nuclear lamina invaginations and DNA damage. DNA stain (white) marks nuclei; YFP (green) marks ISCs; anti-Lamin B (white) marks the nuclear lamina; anti-γH2Av (white) marks DNA damage. (I) Model representing the role of Sdc in *Drosophila* ISCs.

We noted, however, that unlike control ISC nuclei which appeared approximately spherical, with a smooth, round nuclear lamina (Figure 3Ai), the nuclei of Sdc-depleted ISCs presented frequent nuclear lamina invaginations (Figure 3Aiii,C) and aberrant nuclear shapes, with more elongated and lobed nuclei (Figure 3D-E). These phenotypes were also seen with other Sdc RNAi lines and when DIAP1 was co-expressed with Sdc RNAi (Figure S3A), indicating that they were neither an off-target effect of the RNAi nor a secondary effect of apoptotic cell death.

We reasoned that Sdc depletion could impair a relay from the plasma membrane to the nucleus, which would in turn alter nuclear properties. The Linker of Nucleoskeleton and Cytoskeleton (LINC) complex is a conserved multiprotein complex that connects the cytoskeleton with the nucleoskeleton, allowing transmission of mechanical force from the cell surface to the nucleus, regulating nuclear deformation, positioning and functions (Horn, 2014; King, 2023). The *Drosophila* LINC complex consists of two KASH proteins, Klarsicht (Klar) and Msp300, and two SUN proteins, Klaroid (Koi) and the testes-specific SPAG4 (Figure 3B) (Horn, 2014). We hypothesised that Sdc might act via the LINC complex and thus tested whether individual knockdowns of Klar, Koi and Msp300 in ISCs recapitulated the lamina invaginations and nuclear shape changes seen upon Sdc depletion. Only Msp300 knockdown resulted in increased lamina invaginations compared to control nuclei, but not to the same extent as Sdc knockdown (Figures 3C and S3A). Knockdown of Klar or Koi produced only modest nuclear elongation (Figure 3D) and nuclear lobing was unaffected (Figure 3E). This suggests Sdc function is unlikely to be fully accounted for by individual LINC complex proteins, although these proteins may act redundantly (Horn, 2014; King, 2023).

Disruptions to the nuclear lamina are often associated with DNA damage (Gauthier & Comaills, 2021; Graziano et al., 2018). Immunolabelling of DNA double strand breaks with γH2Av (Madigan et al., 2002) revealed striking DNA damage acquisition in ISCs upon Sdc knockdown but not in control ISCs (Figure 3F-G). Increased DNA damage was also seen with a second Sdc RNAi line, albeit at a lower frequency (Figure S3B). These results suggest that Sdc regulates a mechanism protecting the genome. Knockdown of individual LINC complex components from ISCs did not recapitulate the level of DNA damage seen upon Sdc depletion (Figure 3F-G), supporting the notion that Sdc does not mediate its effects via the LINC complex. Notably, DNA damage was not a secondary consequence of apoptosis (Figure S3C), and thus may instead contribute to ISC elimination.

To uncover the relationship between nuclear lamina invaginations and DNA damage, we carefully examined the frequency of both features in a large number of ISCs (Figure 3H). DNA damage was found more frequently in Sdc-depleted ISCs with lamina invaginations compared to those without (Figure 3H), supporting a model whereby the development of nuclear lamina invaginations precedes the acquisition of DNA damage.

Altogether, our results suggest that Sdc regulates nuclear shape and the nuclear lamina, in a manner that is not dependent on single members of the LINC complex. In the absence of Sdc, nuclear aberrations arise early, with lamina invaginations likely preceding acquisition of DNA damage, which then conceivably triggers ISC elimination (Figure 3I).

### Sdc controls nuclear envelope remodeling of neural stem cells and is dispensable from female germline stem cells

Next, to test whether Sdc plays similar roles in other stem cell types, we examined female germline stem cells (fGSCs) and larval neural stem cells (neuroblasts).

In the female *Drosophila* germline (Figure S4A), Sdc depletion from fGSCs did not cause germline loss (Figure S4), suggesting that Sdc is dispensable for fGSC survival. Moreover, we did not detect any evidence of germline stem cysts, which arise upon defective fGSC abscission (Mathieu et al., 2013) (Figure S4A-B), nor DNA damage (Figure S4C), indicating that Sdc is not required for abscission or genome protection in these cells. Our results suggest Sdc is dispensable in fGSCs, supporting previous data (Hayashi et al., 2021).

In the larval *Drosophila* brain, which is highly amenable to live imaging (Cabernard & Doe, 2013; Lerit et al., 2014), neuroblasts undergo rapid asymmetric cell division, generating a large self-renewing neuroblast with a large nucleus and a smaller differentiating ganglion mother cell (GMC) with a small nucleus (Homem & Knoblich, 2012) (Figure 4A). Wildtype neuroblasts undergo semi-closed mitosis, whereby the nuclear envelope persists throughout mitosis, dependent on the maintenance of a supporting nuclear lamina (Roubinet et al., 2021). Some reservoirs of nuclear membrane are also present in the cytoplasm of the renewing neuroblast and contribute to differential nuclear growth of the daughter cells (Roubinet et al., 2021) (Figure 4A-B).

**Figure 4:**
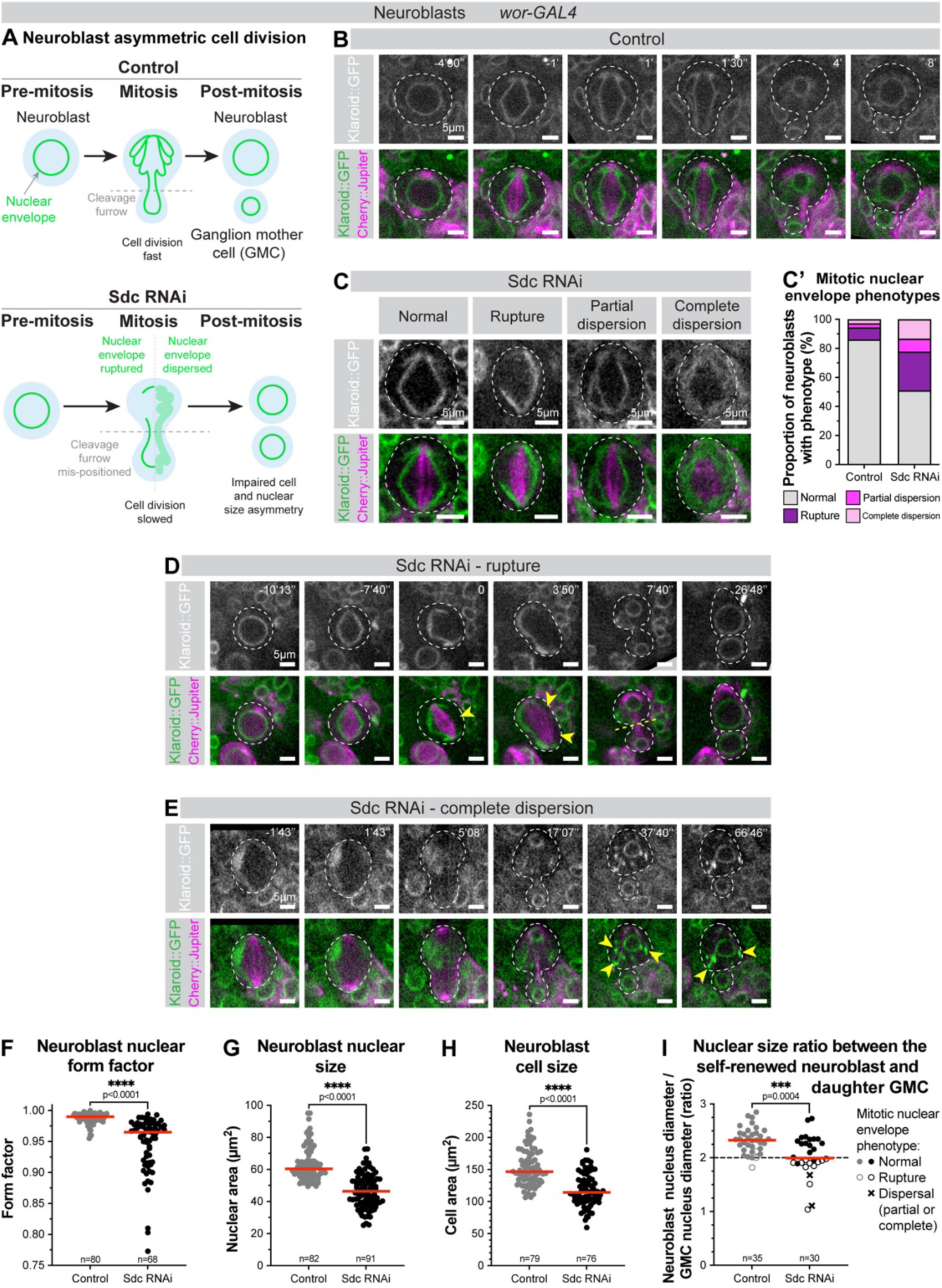
Sdc knockdown in neural stem cells alters nuclear envelope remodeling and asymmetric cell division. (A) *Drosophila* neuroblast division. (B-C & D-E) Klaroid::GFP (white/green) marks inner nuclear membrane; Cherry::Jupiter (magenta) marks mitotic spindle. Neuroblast outlined with white dashed line. Time stamp in minutes and seconds, with anaphase onset defined as time 0. (B) Time lapse of a control mitotic neuroblast. Representative time lapse from 36 neuroblasts from 4 independent experiments (20 brains total). (C) Example images of mitotic neuroblasts expressing Sdc RNAi with mitotic nuclear envelope phenotypes. Representative images from 45 neuroblasts from two independent experiments (7 brains total). (C’) Prevalence of mitotic nuclear envelope phenotypes. n numbers same as B&C. (D) Time lapse of a mitotic neuroblast expressing Sdc RNAi, with a large rupture of the nuclear envelope (yellow arrowheads). Yellow dashed line marks cleavage furrow. (E) Time lapse of a mitotic neuroblast expressing Sdc RNAi, with complete dispersion of the nuclear envelope. Arrowheads mark abnormal nuclear membrane aggregates in the cytoplasm of the renewing neuroblast. (F) Neuroblast nuclear form factor, (G) Neuroblast nuclear size and (H) Neuroblast cell size, measured during interphase. n=number of neuroblast, from 13 brains from 3 independent experiments (control) or 7 brains from 2 independent experiments (Sdc RNAi). (I) Nuclear size ratio between the renewed neuroblast and newly formed GMC, measured at telophase. Neuroblast nuclear envelope phenotype indicated for each data point. n=number of neuroblast/GMC pairs measured, from 13 brains from 3 independent experiments (control) or 7 brains from 2 independent experiments (Sdc RNAi)

After confirming the presence of Sdc in neuroblasts (data not shown), we examined whether its depletion affected cell division and nuclear properties. Sdc-depleted neuroblasts exhibited prolonged cell division (Figure 4B,D-E) and had a variety of cell division, nuclear envelope, and nuclear shape defects. Approximately 50% of mitotic neuroblasts expressing Sdc RNAi displayed abnormal mitotic nuclear envelope remodelling (Figure 4C-C’) including ruptures (Figure 4C-D) and dispersion (Figure 4C,E) of the nuclear envelope, which appeared as early as metaphase. Outside of mitosis, Sdc-depleted neuroblasts had aberrant nuclear shape and size (Figure 4F-G) and abnormal cell size (Figure 4H).

Furthermore, Sdc depletion resulted in abnormal ratio in nuclear size between the renewed neuroblast and the newly formed GMC (Figure 4I). This was sometimes caused by mispositioning of the cleavage furrow and abnormal nuclear membrane partitioning between the daughter cells, resulting in abnormally large GMC nuclei (Figure 4D), and in other cases was caused by impaired nuclear growth, resulting in abnormally small neuroblast nuclei (Figure 4E). These defects in nuclear size ratio were predominantly associated with divisions where the nuclear envelope was either ruptured or completely dispersed (Figure 4I), suggesting that mitotic nuclear envelope remodelling defects induced by Sdc knockdown might have a causative effect on nuclear size ratio.

Thus, in at least two different somatic stem cell types, in the adult gut and developing brain, the transmembrane protein Sdc modulates nuclear properties, with major impacts on cell division and tissue renewal.

## Discussion

Stem cell activity underpins tissue renewal and disease, and stem cells are highly regulated by the complex, dynamic environment in which they reside. The adult *Drosophila* gut has proven an excellent in vivo system allowing functional characterisation of proteins involved in basement membrane adhesion and mechanostransduction, from the molecular to tissue level (Lee et al., 2012; Lin et al., 2013; Okumura et al., 2014; You et al., 2014; Patel et al., 2015; Chen et al., 2018; Howard et al., 2019; Mlih & Karpac, 2022; F. Chen et al., 2021; Ferguson et al., 2021; Bohere et al., 2022). Here we have identified previously unknown connections between the conserved transmembrane proteoglycan Sdc, nuclear properties and stem cell behaviour.

Our temporal analysis of *Drosophila* ISCs depleted of Sdc revealed early nuclear lamina defects (Figure 3A,C) and gradual acquisition of DNA damage (Figure 3G) followed by ISC loss (Figure 1G-H) suggesting that Sdc regulates nuclear lamina organisation and safeguards genomic integrity to protect stem cells. How could Sdc regulate nuclear properties and cell survival? Sdc could prevent initiation of cell death programmes which subsequently alter the nucleus (Ambrosini et al., 2019). However, when we blocked apoptosis nuclear abnormalities were still observed (Figure S3A,C), arguing against such a mechanism.

Alternatively, Sdc could regulate nuclear properties, which in turn promote stem cell survival. In the female *Drosophila* germline, mutations inducing deformation of the nuclear lamina promote DNA damage and stem cell loss (Barton et al., 2018; Duan et al., 2020). This is compatible with our results which show a strong association between nuclear lamina defects and DNA damage in Sdc-depleted ISCs (Figure 3H). However, Sdc did not contribute to fGSC protection (Figure S4), suggesting tissue-specific roles. In ISCs, the proximal cause of death upon Sdc knockdown remains to be determined, although the DNA damage acquired by these cells seems a likely driving force, with physiological DNA damage and genotoxic stress known to contribute to ISC elimination (Nagy et al., 2018; Sharma et al., 2020).

Might Sdc be required at a particular phase of the cell cycle? Sdc depletion in neuroblasts perturbs mitotic nuclear envelope remodelling and impairs cell and nuclear division, with effects seen as early as metaphase (Figure 4A,C-E), revealing additional roles of Sdc in cell division, divergent from those found in mammalian cell culture (Keller-Pinter et al., 2010; Addi et al., 2020). It is notable that in the gut, which is predominantly composed of post-mitotic cells, Sdc depletion was only deleterious in the cell population capable of dividing (Figure 1). Future live imaging of the *Drosophila* gut would allow dynamic assessment of ISC behaviour and analysis of whether nuclear aberrations are precipitated by cell division, as appears to be the case in neural stem cells.

As our results pointed towards a connection between Sdc and nuclear properties, we reasoned that Sdc might be involved in force transmission between the plasma membrane and nucleus (Kalukula et al., 2022; Miroshnikova & Wickström, 2022). Indeed, mammalian Sdc-1 and -4 can act as mechanosensors, transmitting force via their interaction with the cytoskeleton (Chronopoulos et al., 2020; Le et al., 2021), although Sdc’s involvement in mechanical signalling remains understudied in comparison to its involvement in chemical signalling. Excitingly, we found that knockdown of individual components of the LINC complex, stereotypically considered the major machinery relaying force to the nucleus, did not produce similar nuclear aberrations seen upon Sdc depletion, suggesting that Sdc might act via a LINC-independent route. Alternatively, LINC complex components may act redundantly (Horn, 2014), or the environment may influence their requirement. For example, Klar facilitates nuclear movement as ISCs migrate locally during tissue repair but is dispensable under unchallenged conditions when ISCs are relatively immotile (Hu et al., 2021; Marchetti et al., 2022) (Figure 2G).

If changes to the nucleus are not mediated via the LINC complex, we speculate that Sdc regulates the cytoskeleton, with subsequent effects on nuclear properties (Figure 3I). There is a large body of evidence showing that the cytoskeleton can influence nuclear properties (Davidson & Cadot, 2021; Dos Santos & Toseland, 2021; Shokrollahi & Mekhail, 2021; Miroshnikova & Wickström, 2022), and altered cytoskeletal dynamics could also drive cell shape changes induced upon Sdc knockdown (Figure 2B,G). Recent work in mammalian cells has uncovered a laminin-keratin link which shields the nucleus from actomyosin-mediated mechanical deformation, with keratin intermediate filaments forming a protective meshwork around the nucleus (Kechagia et al., 2023). While *Drosophila* cells lack these cytoplasmic intermediate filaments, it is possible that microtubule and/or actin networks instead modulate the nucleus in a force-dependent manner, as reported in other contexts (Almonacid et al., 2019; Biedzinski et al., 2020). Notably, Sdc can directly bind tubulin (K. Chen & Williams, 2013; Gondelaud & Ricard-Blum, 2019) and actin-binding proteins, such as FERM domain-containing proteins (Granés et al., 2003; Gondelaud & Ricard-Blum, 2019). Future work should seek to identify the precise molecular mechanism(s) via which Sdc elicits its effects at the nucleus, and whether these depend on its interaction with the extracellular matrix.

In conclusion, using a simple, tractable *in vivo* model, we have discovered that Sdc regulates nuclear properties and cell function of intestinal and neural stem cells. Whilst there will be microenvironmental differences between cell types and organisms, given Sdc’s high conservation, the broad relationship between Sdc and nuclear properties is likely to be upheld across organisms. This may be therapeutically relevant, especially as nuclear changes and genomic instability are hallmarks of cancer cells, and that Sdc proteins are often deregulated in cancer and inflammatory diseases (Gopal, 2020; Czarnowski, 2021).

## Material and Methods

### Fly strains

All *Drosophila* stocks were maintained at 18°C or amplified at 25°C on standard medium (cornmeal, yeast, glucose, agar, water, Nipagin food medium). Fly strains are referenced in Table 1.

**Table 1.** Fly stocks used in this work.

| Full genotype | Informal name | Reference/source |
| --- | --- | --- |
| w ; esg-GAL4, tub-GAL80ts, UAS-GFP / CyO ; UAS-flp, act>CD2> GAL4 / TM6B ; + / + | *esg<sup>ts</sup>* F/O | (Jiang et al., 2009) |
| + / + ; UAS-CD8-GFP ; + / + ; + / + | UAS-CD8-GFP |  |
| y[1] w[*] ; Mi{Trojan-GAL4.0}Sdc[MI03925-TG4.0] / SM6a ; + / + ; + / + | *syndecan-GAL4* | BDSC 76652 |
| w; esg-GAL4 UAS-YFP / CyO ; Su(H) GBE-GAL80, tub-GAL80ts / TM3, Sb ; + / + | *esgSu(H)* (expresses in ISCs) | Cedric Polesello |
| w; UAS-CD8-GFP, tub-GAL80ts ; Klu-GAL4, UAS-H2B-RFP / SM6a, TM6B ; + / + | *Klu<sup>ts</sup>* (expresses in EBs) | (Reiff et al., 2019) |
| + / + ; UAS-YFP / CDY ; voila-GAL4 / TM6B ; + / + | *pros<sup>ts</sup>* (expresses in pre-EEs & EEs) | Irene Miguel-Aliaga |
| + / + ; Syndecan-Scarlet-2xHA / (CyO) ; + / + ; + / + | Sdc-Scarlet | this study |
| yw ; MyoIA-GAL4, tub-GAL80ts / Cyo ; UAS-GFP / TM6b ; + / + | *MyoIA<sup>ts</sup>* (expresses in ECs) | Kolahgar lab |
| y[1] sc[*] v[1] sev[21] ; P{y[+t7.7] ; + / + ; v[+t1.8]=VALIUM20-mCherry.RNAi}attP2 ; + / + | mCherry / Control RNAi | BDSC 35785 |
| y[1] sc[*] v[1] sev[21] ; P{y[+t7.7] v[+t1.8]=TRiP.HMC03287}attP2 / TM3, Sb[1] Ser[1] ; + / + ; + / + | Sdc RNAi 1 | BDSC 51723 |
| + / + ; UAS-Syndecan RNAi (KK) ; + / + ; + / + | Sdc RNAi 2 | VDRC 107320 |
| + / + ; + / + ; UAS-Syndecan RNAi (GD); + / + | Sdc RNAi 3 (also referred to as Sdc RNAi in main figures) | VDRC 13322 |
| y[1] w[*] ; P{w[+mC]=UAS-Sdc.J}3 ; + / + ; + / + | Sdc-J | BDSC 8564 |
| w[1118] ; +/+ ; +/+ ; +/+ | w<sup>1118</sup> / Control | Celia Garcia Cortes |
| w ; UAS-DIAP1 / Cyo, Kr-GAL4, UAS-GFP ; TM2 / TM6, Df-YFP ; + / + | DIAP1 | Jean-Paul Vincent |
| w ; UAS-DIAP1 / CyO ; UAS-Sdc RNAi (VDRC 13322) / TM6c, Df-YFP ; + / + | Sdc RNAi, DIAP1 | Kolahgar lab |
| + / + ; UAS-AuroraB-RNAi (KK) ; + / + ; + / + | AuroraB RNAi | VDRC 104051 |
| y[1] sc[*] v[1] sev[21] ; P{y[+t7.7] v[+t1.8]=TRiP.HMS02172}attP40 ; + / + ; + / + | Klaroid RNAi | BDSC 40924 |
| y[1] sc[*] v[1] sev[21] ; P{y[+t7.7] ; + / + ; v[+t1.8]=TRiP.HMS01612}attP2 ; + / + | Klarsicht RNAi | BDSC 36721 |
| y[1] sc[*] v[1] sev[21] ; + / + ; P{y[+t7.7] v[+t1.8]=TRiP.HMS00632}attP2 ; + / + | Msp300 RNAi 1 | BDSC 32848 |
| P{UAS-Dcr-2.D}1, w[1118] ; nosP-GAL4-NGT40 | *nos-GAL4* | BDSC 25751 |
| w; Wor-GAL4, KlaroidCB04483, UAS Cherry::Jupiter / Cyo; UAS Dicer / TM6B,Tb | *wor-GAL4* | (Roubinet et al., 2021) |

We used FlyBase (release FB2023_06) to find information on phenotypes/function/stocks/gene expression etc. (Jenkins et al., 2022).

The fly line expressing Sdc-Scarlet-2xHA was produced by Genetivision. The following guide RNAs (gRNA) were selected for CRISPR/Cas9 mediated double strand breaks, for targeting immediately after the final codon of the *syndecan* gene: TATCTCAGGCGTAGAACTCG CGG and GTGTGCGTATGACTGGACGA AGG. The donor template for homology directed repair was designed to contain a four-serine linker, Scarlet and a HA-GG-HA tag.

### Experimental conditions for gut analyses

All analyses were performed on mated females.

For experiments involving GAL80^ts^, flies were reared at 18°C throughout embryonic, larval and pupal development, with crosses flipped every 3-4 days. Flies were collected every 3-4 days after hatching and transferred to 29°C, (which was defined as day 1) and transferred to new food every two days.

For the survival experiment, crosses of 15 *esgSu(H)* virgins and 6 w^1118^ or Sdc RNAi males were set up at 18°C and flipped every 3-4 days. Upon hatching, adult flies were collected and transferred to 29°C for two days to allow mating. On day 2, flies were anesthetised for minimal time to retrieve female flies of the correct genotypes, which were transferred into vials of standard medium at a density of 12 females per vial. All vials were placed horizontally at 29°C, with vial position randomised to control for any variation in temperature or humidity in the incubator. Survival was recorded daily, and flies were transferred to fresh food every 2 days until the end of the experiment, when the final fly died.

*esg^ts^ F/O* (“escargot flip out”) experiments (Jiang et al., 2009) were performed as in (Bohere et al., 2022). At the non-permissive temperature (18°C), GAL80^ts^ which is expressed under the ubiquitous tubulin promoter, inhibits GAL4 preventing expression of UAS-GFP, UAS RNAi and UAS-flp-recombinase. Upon hatching, adult flies are transferred to the permissive temperature (29°C). This inactivates GAL80ts, allowing the intestinal progenitor cell-specific *esg-GAL4* to drive expression of UAS-transgenes. The flp-recombinase excises a STOP codon between the ubiquitous actin promotor and GAL4, resulting in permanent and heritable GAL4, and thus UAS-transgene expression, regardless of cell type. Thus, all cells which arise from progenitors after the temperature shift will express GFP, providing a visual readout for new cell production.

### Gut dissection and immunostaining

Midguts were dissected in 1X PBS (Oxoid, BR0014G) and fixed for 20 minutes at room temperature in fresh 4% methanol-free paraformaldehyde (Polysciences, 18814-10) diluted in PBS, 0.025% Triton X-100 (Sigma Aldrich, X100). Samples were given three 5-minute washes in 0.25% PBST and permeabilised for 30 minutes in 1% PBST. Samples were then incubated for 30 minutes at room temperature in blocking buffer (0.1% PBST, 0.1% BSA (Sigma, A2153-10G)), followed by incubation overnight at 4°C with primary antibodies diluted in blocking buffer. Samples were next given three 20-minute washes in 0.25% PBST and incubated for 2 hours at room temperature in secondary antibodies diluted in 0.25% PBST. Finally, guts were given three 20-minute washes in 0.25% PBST. Guts were mounted in Vectashield (Vector Laboratories, H-1000) on a glass slide (VWR International, SuperFrost 1.0mm, ISC 8037/1) with coverslip (Menzel-Gläser, 22×50mm#1, 12342118). The following primary antibodies were used: chicken anti-GFP, 1/1000 (Abcam, ab13970); rat anti-α-catenin, 1/20 (DSHB, D-CAT-1-s); rabbit anti-HA, 1/500 (Cell Signaling, 3724T); rabbit anti-H2Av, 1/700 (Rockland, 600-401-914); mouse anti-LaminDm0 (Lamin B), 1/10 (DSHB, ADL84.12). Alexa-488-, 555-, and 647-conjugated secondary goat antibodies (Molecular Probes) were used. F-actin was stained with Phalloidin (Molecular Probes, A12380 or A22287). Nuclei were stained with DAPI (Molecular Probes, D1306) or Hoechst (Molecular Probes, H1399).

### Ovary dissection and immunostaining

Ovaries were dissected and collected in 1X PBS and then fixed for 25 minutes in 4% formaldehyde in 0.3 PBSTX (0.3% Triton-X in 1X PBS) at room temperature. Samples were given three 15-minute washes in 0.3% PBSTX and then incubated for one hour in blocking buffer (0.2 ug/μl BSA in 0.3% PBSTX), followed by incubation overnight at 4°C with primary antibodies and Phalloidin diluted in blocking buffer. Samples were then given three 15-minute washes in 0.3% PBSTX and incubated for 2 hours at room temperature in secondary antibodies diluted in blocking buffer. Samples were then given three 15-minute washes in 0.3% PBSTX, including Hoechst (33342) DNA stain in the first wash. Samples were mounted in VectaShield media (Vector Laboratories, H-1000). The following primary antibodies were used: mouse anti-H2Av, 1/200 (DSHB UNC93-5.2.1); mouse anti-α-Spectrin, 1/200 (DSHB, 3A9).

### Image acquisition and processing

Confocal images of fixed samples were acquired on a Leica TCS SP8 advanced confocal microscope with laser gain and power consistent within experiments. Gut overview images were acquired on a Leica M165FC microscope with Leica DFC3000G camera or EVOS M7000 (ThermoFisher Scientific). All images were processed and analysed with FiJi (Schindelin et al., 2012). Imaris 3 64 7.5.2 was used to analyse neuroblast data presented in Figure 4. Figures were compiled in Adobe Illustrator.

Unless otherwise specified, all images show the R4/5 posterior midgut region. Midgut surface views are z-projections through half the gut depth. Images of single ISCs for nuclear lamina and shape qualifications are single z sections.

### Live imaging of larval neuroblasts

Live imaging experiments were performed on intact brains. Larvae expressing nuclear membrane and spindle marker (Klaroid::GFP and UAS-Cherry::Jupiter) together with UAS-Dicer, UAS-Sdc RNAi (VDRC 13322) and *worniu-GAL4* were dissected seventy two or ninety-six hours after egg laying, respectively, in imaging medium (Schneider’s insect medium mixed with 10% fetal bovine serum (FBS) (Sigma, F7524), 2% PenStrepNeo (Sigma, P4083), 0.02 mg/mL insulin (Sigma, 11070), 20mM L-glutamine (Sigma, G8540), 0.04 mg/mL L-glutathione (Sigma, G4251) and 5 mg/mL 20-hydroxyecdysone (Sigma, H5142)) warmed up to room temperature before use. Brains were then transferred onto IbiTreat micro-slide 15 well 3D (Ibidi, 81506) and imaged with a confocal spinning disc. All images presented in Figure 4 are single z sections. Anaphase onset is defined as t=0.

### Live imaging and analysis of adult intestinal stem cells

Guts expressing YFP +/− UAS Sdc RNAi in ISCs (with *esgSu(H)* system) were carefully dissected 7 to 16 days after induction of GAL4 expression at 29°C in Shield and Sang M3 medium (Sigma S3652), keeping the crop, Malpighian tubules and ovaries attached. Guts were mounted in imaging medium (Shields and Shang M3 medium, supplemented with 2% Fetal Bovine Serum, 0.5% penicillin-streptomycin (Invitrogen, 15140–122) and methylcellulose (Sigma, M0387-100G) 2.5% wt/vol, to stabilise the gut in the chamber (Aldaz et al., 2010) between a concanavalin-A coated coverslip and an oxygen permeable membrane and left to settle for 5min. All guts were live imaged for 10 to 15 hours with a 2-minute interval between each scan comprising 20 to 25 sections of 1μm, on a 40X air objective using the EVOS M7000 (ThermoFisher Scientific). Image files were processed in FiJi, using Bleach correction and Stackreg plugins. Regions of interest (25×25μm) were cropped, upscaled to 512×512 pixels and reduced to 264 images (528min). Cells shapes at specified timepoints were drawn and superposed using Adobe Illustrator.

### Image analysis and quantifications

#### Cell density and cell type proportion

The total number of epithelial cells (using nuclei as a proxy) and the total number of each cell type (identified by their expression of fluorescent protein or nuclear size (small, medium and large cells in *esg^ts^ F/O* experiment)) were manually counted using the cell counter tool in Fiji, and a ratio was calculated. Cell density was calculated as the total number of cells per 80×200μm area. Measurements were done on z projections of half the intestinal epithelium depth, except for Figure 1E-F and Figure S1B, where single z slices were analysed, on both sides of the gut tube where possible.

#### Posterior midgut progenitor cell retention and new cell production

Posterior midguts were visually inspected and manually assigned to one of five phenotypic categories.

#### Intestinal stem cell size and shape

Masks of ISCs were manually drawn around cytoplasmic YFP signal with the Fiji freehand line on z-projections encompassing half the gut depth. The following parameters were calculated with the Fiji ‘analyse particles’ function: size, aspect ratio (defined as the major axis / minor axis, where a value of 1.0 indicates a shape with equal dimensions in both axes, and higher values indicate increasingly elongated shapes) and form factor (defined as 4π x area / perimeter^2^, where a value of 1.0 indicates a perfect circle and lower values indicate increasingly convoluted shapes).

#### Nuclear lamina invaginations and nuclear shape

High magnification single z confocal images of individual ISCs were acquired through the centre of the nucleus and processed with the Cell Profiler pipeline described in (Janssen et al., 2022). Information was extracted about nuclear eccentricity (defined as c / a, where c^2^ = a^2^ – b^2^, with a defined as half the length of the equivalent eclipse and b as half the width of the equivalent eclipse. A resulting value of 0 indicates a perfect circle and higher values indicate increasingly elongated shapes) and nuclear form factor (defined as 4π x area / perimeter^2^, where a value of 1.0 indicates a perfect circle and lower values indicate increasingly convoluted shapes). To identify Lamin B which localises to the nuclear periphery versus that which forms invaginations, the nucleus is shrunk by a defined number of pixels. High Lamin B areas located outside this shrunk nucleus are defined as corresponding to the nuclear periphery, whereas high Lamin B areas located within the shrunk nucleus, in the nuclear interior, are defined as corresponding to invaginations.

#### DNA damage

ISCs were scored DNA damage positive if they possessed a bright, large H2Av puncta colocalising with the nucleus or H2Av staining covering a large amount of the nuclear area. The proportion of ISCs with DNA damage was calculated for each gut as the number of ISCs with DNA damage divided by the total number of ISCs.

#### Neuroblast nuclear and cell size

Areas were obtained by manually drawing around the border of each nucleus and cell using the Fiji freehand line tool. For quantification of nuclear division asymmetry, the nuclear diameter corresponding to the daughter neuroblast nucleus and to the daughter ganglion mother cell nucleus were measured using ImageJ, after completion of cytokinesis once nuclear growth is finished.

### Statistical analyses

Data were plotted with GraphPad Prism 10 for Mac OS X. In all graphs, red line represents median. For pairwise comparisons, statistical significance was calculated using a Mann–Whitney test and p values <0.05 were considered statistically significant. Categorical data were compared using the Chi square test. Survival was assessed using the Log-rank test.

## Abbreviations

Sdc: Syndecan
ISCs: intestinal stem cells
LINC: Linker of Nucleoskeleton and Cytoskeleton
Koi: Klaroid
Klar: Klarsicht
DIAP1: *Drosophila* inhibitor of apoptosis
EB: enteroblast
pre-EE: pre- enteroendocrine cell
EE: enteroendocrine cell
EC: enterocyte
fGSC: female germline stem cell
GMC: ganglion mother cell
*esg*: *escargot*
*Klu*: *Klumpfuss*
*Myo1A*: *Myosin 1A*
*pros*: *prospero*
*wor*: *worniu*
*nos*: *nanos*

## Acknowledgements

We thank colleagues listed in the Materials & Methods section; the Bloomington *Drosophila* Stock Center; the Vienna *Drosophila* Resource Center and the Developmental Studies Hybridoma Bank for reagents, and the Cambridge Advanced Imaging Facility for use of the confocal facility. We thank the Department of Genetics, University of Cambridge, Fly Facility. We thank Celia Garcia-Cortes for help with ISC live imaging; Nick Brown and Sonali Sengupta for feedback on the manuscript; and Roland Le Borgne and Buzz Baum for access to facilities for larval brain experiments, advice to C.R and for funding: Agence Nationale de la Recherche (Le Borgne Lab: ANR-20-CE13-0015) and Cancer Research UK Discovery Programme (Baum Lab: Grant Award 28276). This work was supported by a Wellcome Trust and Royal Society Sir Henry Dale Fellowship to GK (206208/Z/17/Z), a Wellcome Trust PhD studentship to BLET (102175/B/13/Z), funding from the School of Biological Sciences at the University of Cambridge to JGB, a Herchel Smith Postdoctoral Fellowship to TJS, a Wellcome Trust and Royal Society Sir Henry Dale Fellowship (206257/Z/17/Z) and a Human Frontier Science Program (CDA-00032/2018) to FKT. CR acknowledges funding from the European Union’s Horizon 2020 Research and Innovation Programme under the Marie Skłodowska-Curie (Grant Agreement No 899546).

## Open Access

For the purpose of Open Access, the authors have applied a CC BY public copyright licence to any Author Accepted Manuscript (AAM) version arising from this submission.

## Author contributions

Conceptualisation: BLET and GK; Methodology: BLET, JGB, CR; Formal analysis: BLET, JGB, CR, AB, TJS; Investigation: BLET, JGB, CR, AB, TJS; Writing – Original Draft, BLET and GK; Writing – Review & Editing, all authors; Funding Acquisition, BLET and GK; Supervision, GK

## Declarations of interest

The authors declare no competing interests.

## Supplemental Figures and Legends

**Figure S1:**
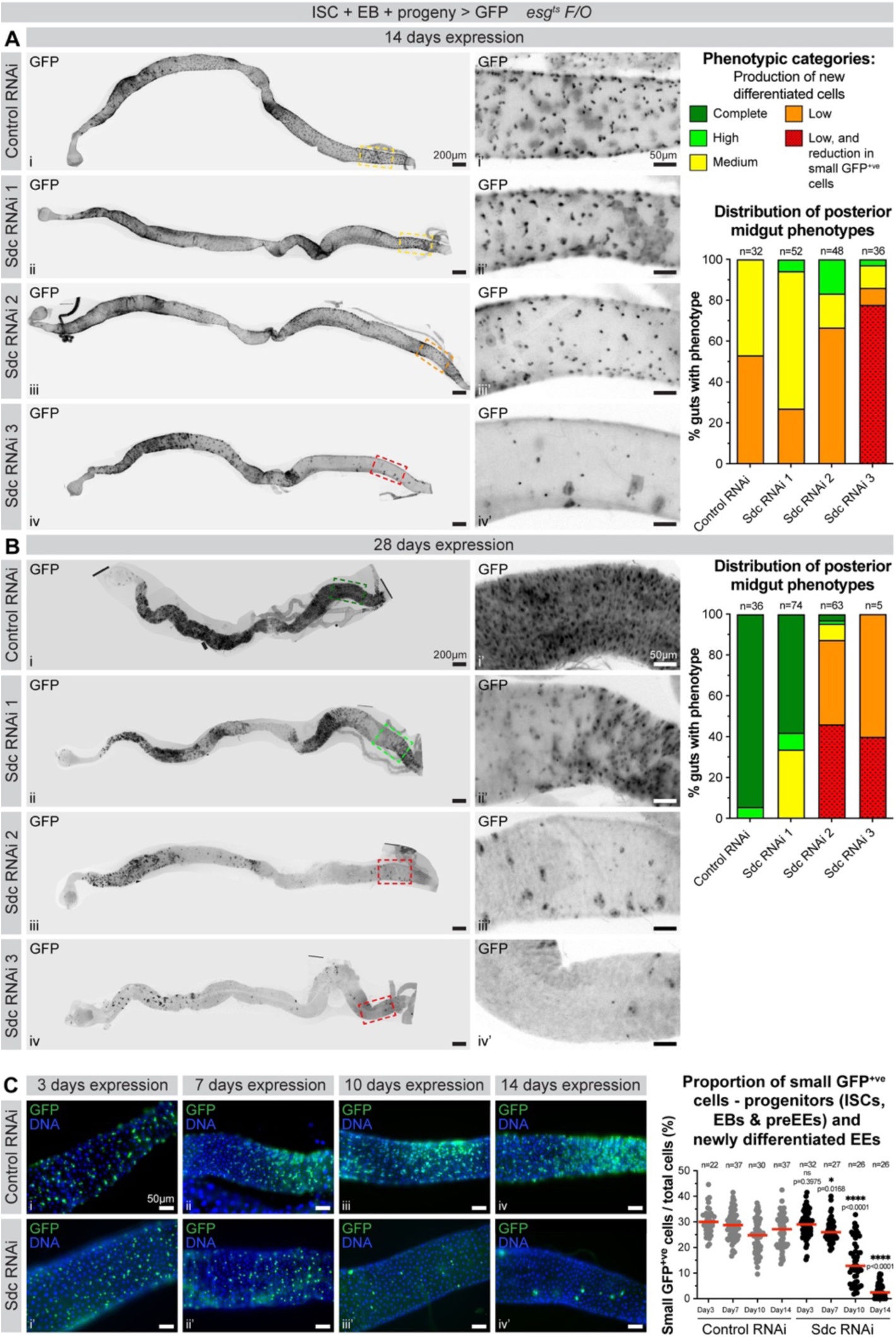
Sdc knockdown in progenitor cells, using three independent RNAi lines, causes progressive progenitor cell loss and failure in new differentiated cell production. (A & B) Whole midgut views (i-iv) and posterior midgut zooms (i’-iv’) from flies expressing control RNAi (i) or one of three Sdc RNAi lines (ii-iv) using the *esg^ts^ F/O* system. Anterior left, posterior right. GFP (black) marks progenitor cells and their progeny. Dashed boxes indicate zoomed area, with the colour of the box indicating the phenotypic category to which the gut was assigned. Graphs show the distribution of posterior midgut phenotypes, with the size of the coloured bar representing the proportion of guts assigned to the phenotypic category in each genotype. n=number of guts, from three replicates. (C) Surface views of midguts expressing control RNAi (i-iv) or Sdc RNAi (i’-iv’) using the *esg^ts^ F/O* system. DNA stain (blue) marks nuclei; GFP (green) marks progenitor cells and their progeny. Graph shows the proportion of small GFP^+ve^ cells (corresponding to progenitors (ISCs, EBs and pre-EEs) and newly differentiated EEs). n=number of guts, from three replicates.

**Figure S2:**
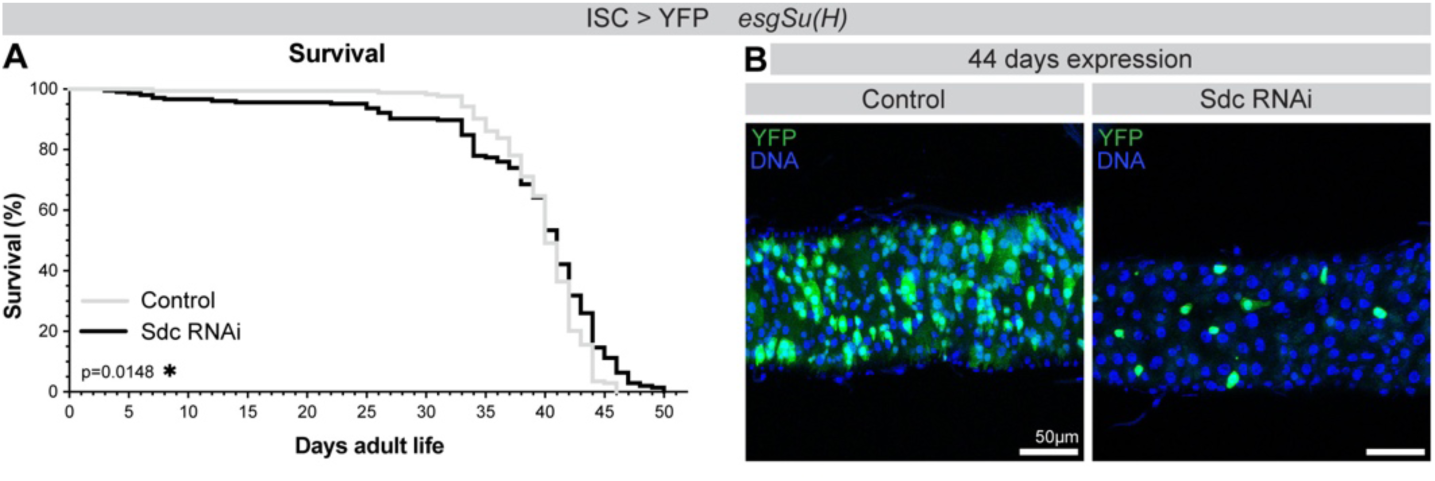
Sdc knockdown in ISCs does not compromise fly survival under unchallenged conditions. (A) Survival during continuous feeding (unchallenged conditions). Log-rank test. (B) Surface views of control midguts, or midguts expressing Sdc RNAi using the *esgSu(H)* system. DNA stain (blue) marks nuclei of all cells; YFP (green) marks ISCs.

**Figure S3:**
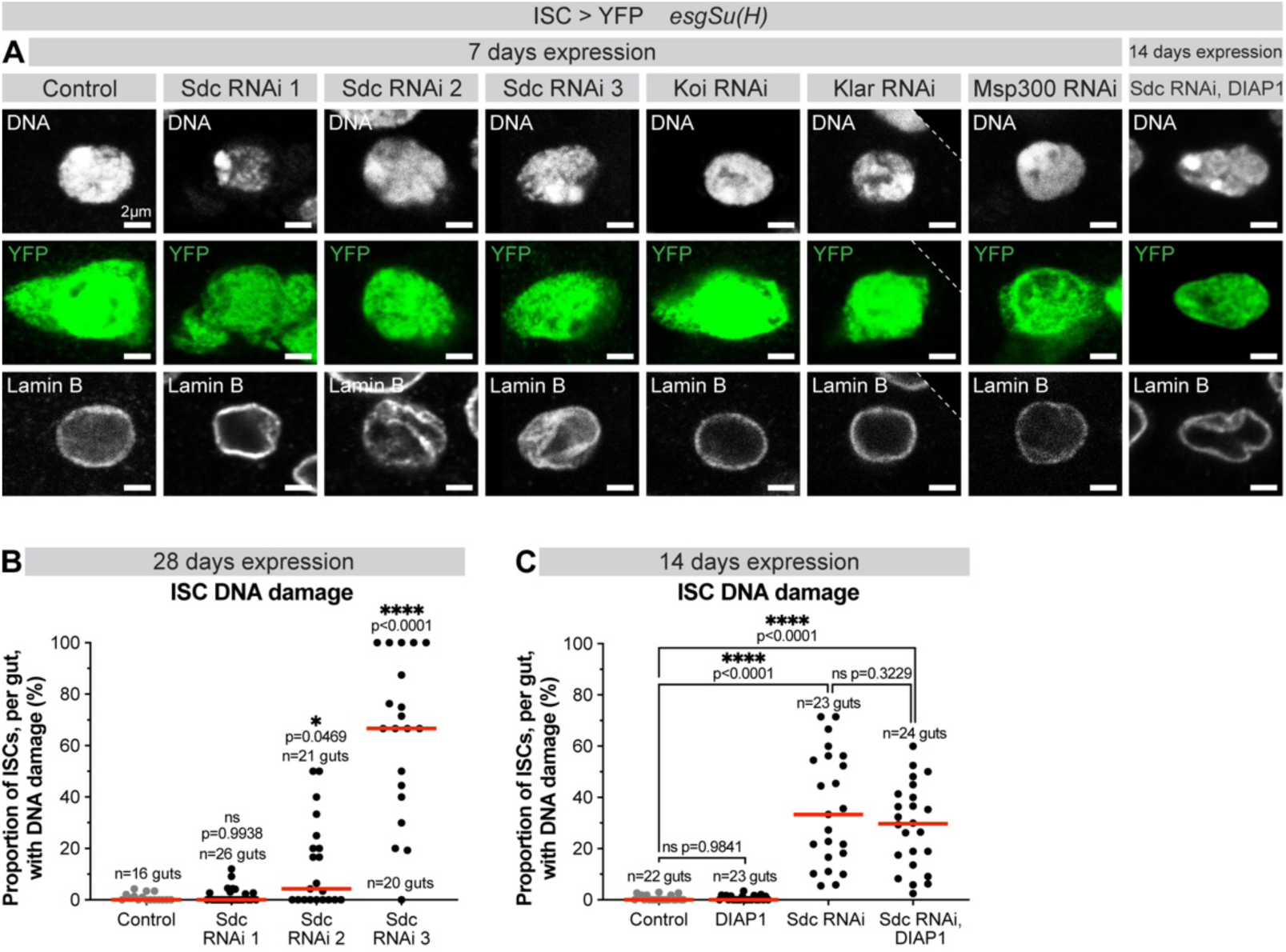
Sdc knockdown, but not knockdown of LINC complex components, in ISCs causes these cells to acquire nuclear lamina invaginations and DNA damage. (A) Control ISC, and ISCs expressing various RNAis. DNA stain (white) marks nuclei; YFP (green) marks ISCs; anti-Lamin B (white) marks nuclear lamina. (B) Proportion of ISCs, per gut, with DNA damage. n=number of guts, from three replicates. >100 ISCs analysed per genotype. (C) Proportion of ISCs, per gut, with DNA damage. n=number of guts, from three replicates. >400 ISCs analysed per genotype.

**Figure S4:**
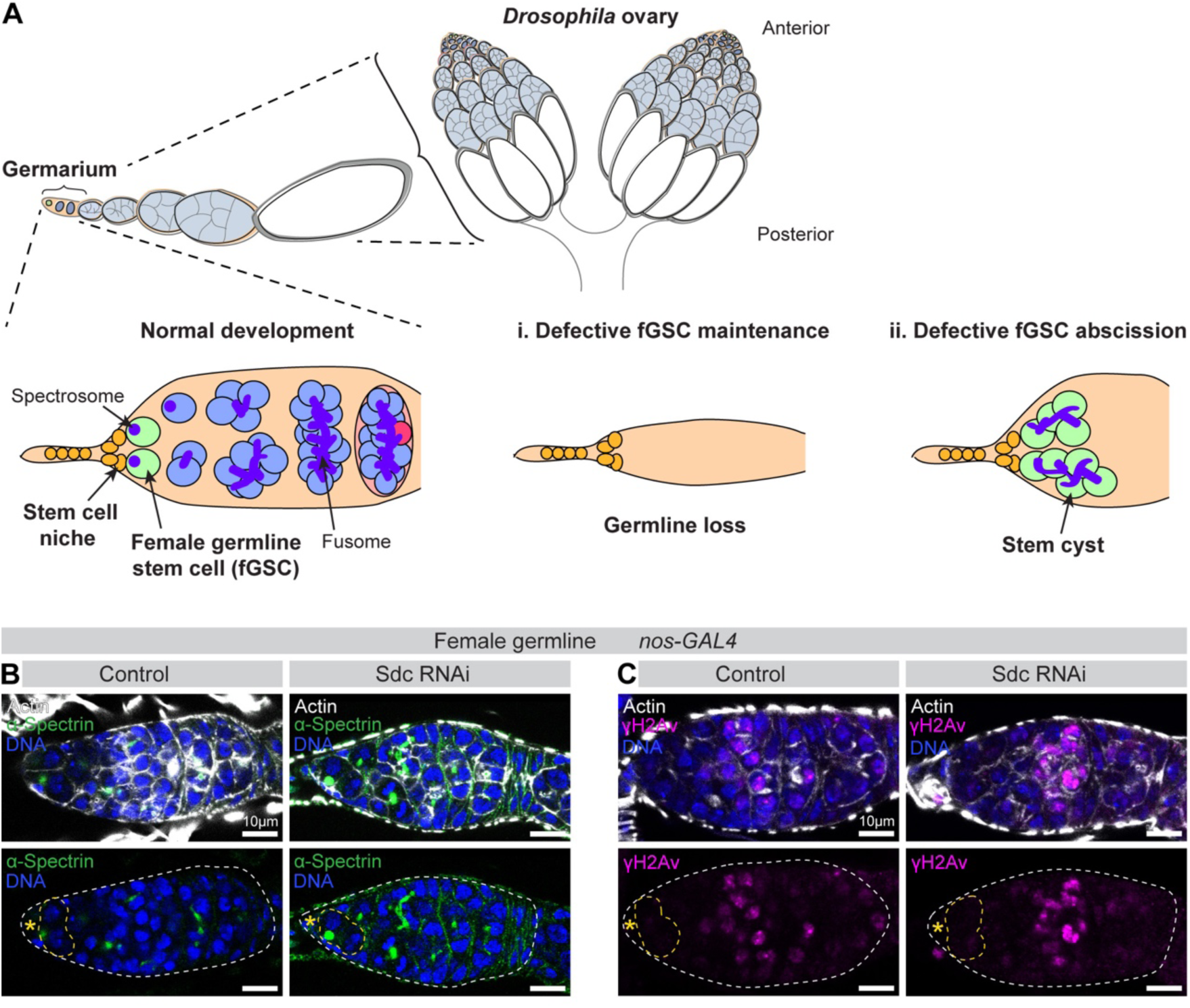
Sdc is dispensable from female germline stem cells. (A) Schematic of female germline development. Female germline stem cells (fGSCs) are maintained in a stem cell niche at the anterior of the ovary in a structure called the germarium. fGSCs provide an excellent model for identifying factors involved in stem cell maintenance (i) and abscission (ii), with clear phenotypic readouts (Sanchez et al., 2016). (B) Germaria from control and germline-specific knockdown of Sdc. White dashed line outlines germarium, yellow dashed line outlines fGSCs, yellow asterisk indicates stem cell niche. DNA stain (blue) marks nuclei; Phalloidin (white) marks F-actin; anti-α-Spectrin (green) marks spectrosome/fusome, allowing identification of fGSCs. (C) Germaria from control and germline-specific knockdown of Sdc. White dashed line outlines germaria, yellow dashed line outlines fGSCs, yellow asterisk indicates stem cell niche. DNA stain (blue) marks nuclei; Phalloidin (white) marks F-actin; anti-γH2Av (magenta) marks DNA damage.

## Notes

### Competing Interest Statement

The authors have declared no competing interest.

